# Spatial filters – In search of the virtual electrode

**DOI:** 10.1101/2023.09.19.558390

**Authors:** Alain de Cheveigné

## Abstract

A spatial filter is a set of weights which, applied to multichannel data, creates a “virtual channel” with desirable properties. Examples range from simple hard-wired spatial filters (re-referencing, gradient, Laplacian, etc.) to more complex data-driven transforms (beamforming, independent component analysis, etc.). The principle is straightforward, but the properties are obscured by the high dimensionality of the data, and the various “spaces” in which a “spatial” filter operates, inhabited by sources, sensors, fields, or signals. This paper reviews the properties and limits of spatial filters, with particular focus on the popular concept of a “virtual electrode” created by combining channels of a non-invasive recording technique such as EEG or MEG. Whereas a spatial filter can perfectly suppress one or more sources (as many as there are sensors, minus one), it cannot select one source and suppress all others, as might a real electrode. This puts hard limits on what to expect of a virtual electrode, and suggests a slightly different perspective, that of a “virtual scalpel”. Spatial filtering, like temporal filtering, plays an important role in brain data analysis.

**Significance statement:** Spatial filters are ubiquitous in brain data analysis. This paper reviews their properties and the limits of what can be achieved, with emphasis on aspects that are obscured by the complex geometry of sources and sensors, and the high dimensionality of the data.

## Introduction

To understand the brain, we can observe its *electrical activity*, in complement with other observations (e.g. optical) and under the guidance of prior knowledge and theory. However, that goal is thwarted by at least three factors that distort the observations and confuse our understanding. A first factor is that there may be many sources of interest that we need to observe concurrently to understand the neural computations. This is addressed in part by multichannel electrode arrays (e.g. Van Beest et al 2025), or optical imaging techniques.

A second factor is that each electrode or sensor picks up activity of *multiple sources* (Fig. 1A). For invasive recordings (in animal models or occasionally in surgical patients), electrodes may be inserted into the neural tissue, each close to a target source, limiting sensitivity to spurious activity. However, for non-invasive methods such as electroencephalography (EEG) or magnetoencephalography (MEG) the problem is acute. Multichannel recording techniques are useful here too, in that they may allow unwanted sources to be suppressed or attenuated by applying a *spatial filter* to the data (Fig. 1B), an idea captured by the popular term “virtual electrode”. A third factor is the very large number of sources involved, that may limit the effectiveness of such processing. These factors interact in complex fashion that this paper seeks to clarify. It is worth noting that multichannel techniques play two distinct roles in this context: to observe multiple target sources (and possibly locate them in space), and to attenuate unwanted sources.

A spatial filter is a vector of weights applied to multichannel data to obtain a signal (time series) with desirable properties (Fig. 1B). Spatial filters include *hard-wired* transforms such as re-referencing or gradients, and *data-driven* transforms such as principal component analysis (PCA), beamforming or independent component analysis (ICA). The purpose is typically to enhance one or more target sources, or suppress one or more interfering sources, or linearly transform the data for some purpose such as decoding, or a brain-computer interface (BCI), or spike sorting for multichannel electrode arrays, or image enhancement for optical techniques. The term is sometimes also applied to a set of filters organized as matrix, each column of which is a filter in the previous sense (Fig. 1C). Thus, “spatial filtering” can result in a single time series (as from a “virtual electrode”) or a matrix of concurrent time series which alternatively can be seen as spatially-filtered “snapshots” of neural activity patterns over space. Which sense is intended should be clear from the context.

The concept is simple (a filter is a vector of numbers) but it may be hard to grasp the benefits and limits of what can be achieved. “Spatial” might refer to the 3D space of source or sensor positions (5D if orientations are included), or to the distribution of the electric or magnetic field within the brain (for example decomposed into spatial harmonics), or to the *J*-dimensional signal space spanned by the *J* channels of data. The complex geometry of sources within the brain, and the high dimensionality of multichannel signals, conspire to make it hard to understand what is going on, what can be achieved, and whether it is worth counting on new methods to bring improvements over what we have today.

The ambitions of spatial filtering are epitomized by the popular concept of *virtual electrode*, which refers to a spatial filter designed to produce a time series similar – ideally – to that recorded by a real electrode inserted within the brain (Fig. 2). Like a real electrode, the filter targets a particular *spatial position*, and produces a single *time series*. By adjusting its coefficients, a virtual electrode is positioned – virtually – anywhere within the brain, with a spatial pattern of gain that mimics – we hope –attenuation with distance from the tip of a real electrode. The prospect of clinical applications is appealing and constitutes a drive to refine the methods and extend their applicability. A hospital administrator who hesitates at the cost of installing and running a MEG system might be swayed by the perspective of saving time, cost and risk to patients, over surgical implantation of deep electrodes. It is important for him or her (and for the patients) to know whether a virtual electrode can indeed replace a real electrode. Likewise, a cognitive scientist might dream of probing the activity of specific locations within the brain by means of a virtual electrode. It is important for him or her (and whoever reads their results) to understand to what degree the outcome of such “virtual probing” can be trusted.

The aim of this paper is to facilitate this understanding, underscoring both the power of spatial filtering and its limits.

### A diversity of filters

Examples of filters are shown in Fig. 1C. From left to right: the first filter forms a sum of channels, selected perhaps because they are expected to all respond to an underlying source, for example channels over auditory cortex in an experiment involving auditory stimulation. The next filters (organized as a matrix) calculate differences between pairs of channels, as in the gradiometers of an MEG machine. The next subtract the third channel from all channels (re-referencing, common in EEG), The next calculate the second-order gradient (often referred to as current source density, CSD, Wójcik 2022) along a linear electrode array, The next apply columns of a PCA transform matrix, and the final, rightmost filter matrix applies a LCMV (linear constrained minimum variance) beamformer to each of 100 locations in a source space. The first four filter matrices are pre-defined (hard-wired), the last two are the result of a data-driven analysis algorithm. Other examples of hardwired filters are signal space separation (SSS, Taulu and Simola 2006) or synthetic gradiometers (Vrba and Robinson 2001).

**Fig. 1.**
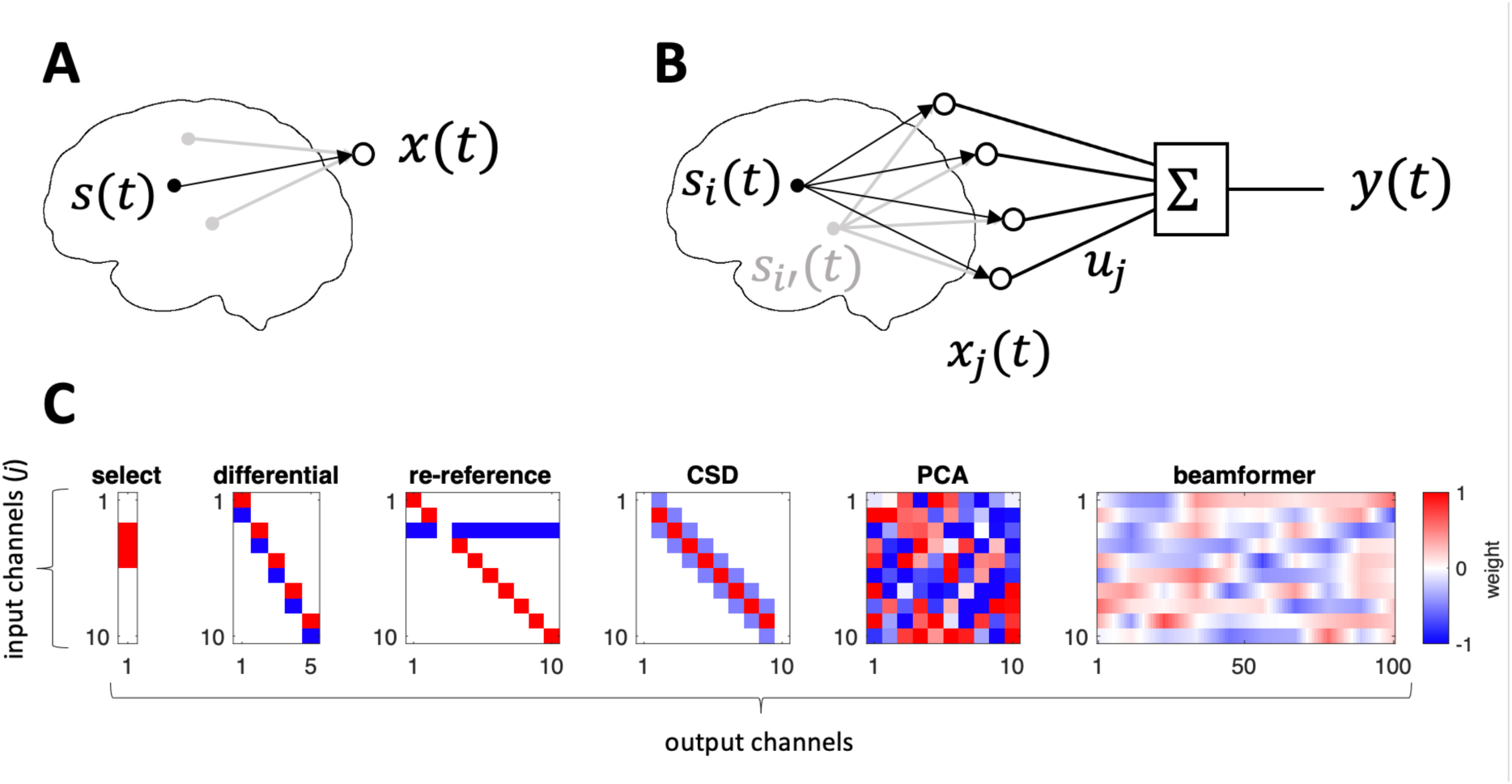
A: An electrode or sensor picks up a source of interest, but the signal is contaminated by other sources. B: Signals from an array of sensors are combined linearly (spatial filter). C: Examples of matrices of spatial filters (each column is a different spatial filter). From left to right: data are summed over selected channels, sensor channels are grouped by pairs into differential channels, channels are re-referenced by subtracting channel 3, channels are combined to estimate the second order gradient (sometimes referred to as current source density, CSD), data are weighted by a PCA matrix, data are weighted by a beamformer matrix (each column targets a different spatial location).

#### What is achieved?

The filter might *enhance* specific activity of interest, or *suppress* specific interference, or merely rearrange the data, for example to reduce their dimensionality, orthogonalize the matrix of time series (remove correlations between channels), or spatially whiten them. Differential and CSD filters enhance local gradients, and they and re-referencing are useful to remove common-mode interference. PCA concentrates data variance in the first few components, and beamforming implements a *virtual electrode* as it is moved through source space (each column targets a different location). A spatial filter does not affect the *spectral* content of the data, other than by giving different weights to channels with different spectral contents, nor does it remove a *DC bias*, nor does it implement *non-linear transforms* as might arise within a neural network for the purpose of classification.

More particularly, a virtual electrode (or “virtual sensor”) is a spatial filter that aims to reproduce the spatial selectivity of a real electrode inserted into neural tissue (Fig. 2). Some empirical success has been reported, for example in recovering the position and time course of sources picked up with real electrodes in epileptic patients undergoing pre-surgical recording with stereotactic EEG (sEEG) electrodes inserted deep in the brain (e.g. Juárez-Martinez et al 2018; Alberto et al 2021; Cao et al 2022, Velmurugan et al 2022; Coelli et al 2023).

**Fig. 2.**
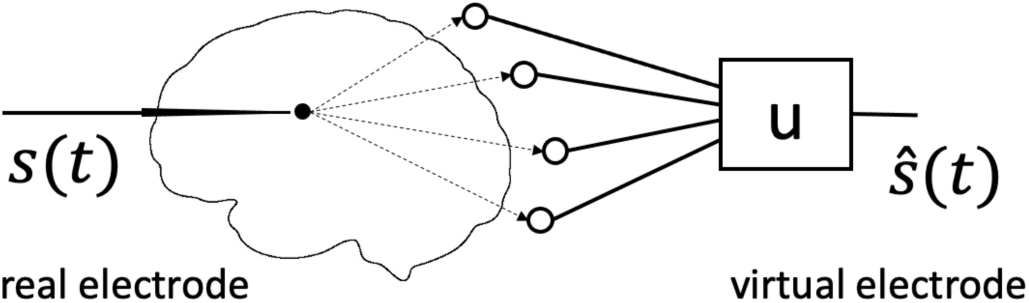
A real electrode (left) is selectively sensitive to brain sources near its tip, and blind to distant sources. A virtual electrode (right) attempts to reproduce this property by applying a spatial filter **u** to the multichannel observation.

A simple mathematical argument tells us that a virtual electrode is not the same as a real electrode. A real electrode placed at one location picks up, potentially, activity different from any other location. If an array of such electrodes were placed in *I* distinct locations (e.g. each of the ∼120000 cubic millimeters of the human brain), the rank of the data matrix could potentially reach *I*. In contrast, the rank of the data matrix obtained by emulating this array with *I* virtual electrodes cannot exceed the number *J* of sensors or electrodes. Since *J* ≪ *I*, these data matrices are necessarily different, and thus virtual electrodes cannot be the same as real electrodes.

A similar argument can be made with respect to electromagnetic (EM) source imaging. At any point in time, only *J* measured values are available to construct the *I* pixels or voxels of the “image” and, likewise, the time course of any pixel or voxel is a function of just *J* sensor signals, in contrast to the *I* ≫ *J* time courses – potentially independent – of the pixels or voxels of an optical imaging technique.

These limitations are, of course well known: they reflect the fundamental non-uniqueness of the inverse problem (Baillet 2010). As in other scientific fields – and indeed perception itself (Helmholtz 1867) – additional assumptions are required in the form of an internal model constrained by the observations to allow *inference* of source activity or locations. This works well, witness the success of EM (electromagnetic) imaging methods in neuroscience, nevertheless, we may (indeed we should) feel uneasy using the outcome of a model-based inference process as *evidence*, because it is unclear to what degree that evidence reflects the observed data or the priors included in the model (Tarantola 2006). Is the sharp boundary of activity, apparent in the image, a reflection of the data, or of the prior? Relatedly, are the imperfections that we observe due to *method-related* or *equipment-related* limitations (amenable to progress), or *fundamental* constraints (not amenable to progress). A goal of this paper is to help answer such questions or, at least, formulate them.

### Some theory

The aim of this section is to point out some basic properties of spatial filters and gain an intuitive understanding of what they can and cannot do. The same ideas are illustrated visually in the next section; the busy reader might want to skip forward and come back for reference.

Recorded data take the form of a matrix **X** of size *T* × *J* where *T* is the number of samples and *J* the number of channels (electrodes or sensors). The recorded data are assumed to reflect the activity of *I* sources within the brain via a linear mixing process:

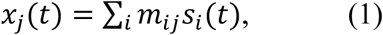

in matrix notation **X** = **SM**, where **S** of size *T* × *I* represents the activity of the sources and **M** is a mixing matrix of size *I* × *J*. Noise sources are included in **S**, hence no noise term in Eq. 1. The matrix **X** is observed, **S** and **M** can only be inferred.

We are usually interested in the activity of certain brain sources, i.e., one or more columns of **S**. Since the sources were mixed linearly, it makes sense to try to recover them using linear methods. Typically, an analysis matrix **U** of size *J* × *K* is applied to the data to yield a matrix **Y** of size *T* × *K* of transformed data:

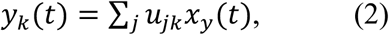

in matrix notation **Y** = **XU**. Each column of **Y** (a time series) is referred to here as a *component*. *K* can be smaller or larger than *J*, however if it is larger the components are linearly dependent and thus correlated: the rank of **Y** cannot be greater than that of **X**.

Ideally, we would like the analysis matrix **U** to be an “unmixing matrix” so that **MU**=**I**, the identity matrix. Each column of **Y** would then map to a source. Unfortunately, brain sources number in billions, whereas sensor channels range from tens to thousands (*I* ≫ *J*), so **M** is not invertible and therefore recovering **S** is impossible. We must settle for the less ambitious objective of finding a transform that produces components that are somehow more *informative* than the raw data. For example, a column of **Y** might approximate the activity of a particular source, or the compound activity of several sources with a functional meaning such as an event-related potential (ERP). Often, we are interested only in one such time series, and thus in just one column **u** of the analysis matrix which constitutes a *spatial filter*, a vector of weights of size *J* which, applied to the observations, produces a single, univariate time series.

#### Vector space

It is helpful to see neural activity, or recorded data, as belonging to a *vector space*, because concepts such as *subspace*, *span*, *dimensionality*, *linear dependence* and *orthogonality* allow for convenient short-cuts and useful insights. The brain source signals *s_i_*(*t*) span a *signal space* of dimension *I* at most (it might be smaller if some of the *s_i_*(*t*) are linearly dependent). The observed data *x_j_*(*t*) span a *subspace* of that space, of dimension at most *J*. The mixing matrix **M** projects from the source space onto this subspace. Since *I* ≫ *J*, a given observation *x_j_*(*t*) might reflect many different patterns of brain activity, a manifestation of the non-uniqueness mentioned earlier. This is schematized in Fig. 3, which shows how two different brain sources might give rise to the same observation when projected on to the lower-dimension observation space.

**Fig. 3.**
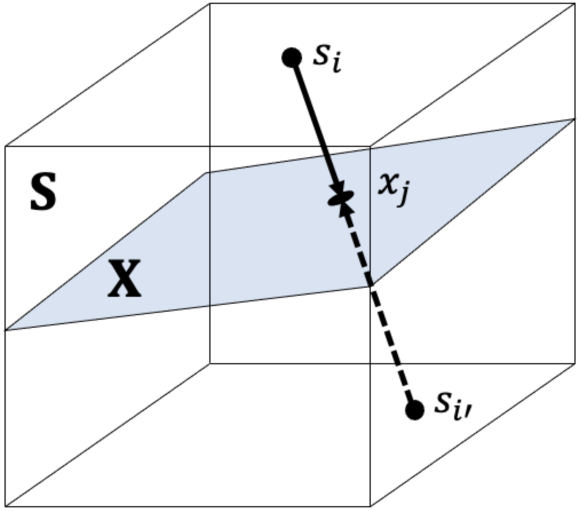
The space spanned by the columns of the observation matrix **X** is a subspace of the space spanned by the brain signals **S**. Brain source signals are projected onto this subspace, and thus multiple sources may map onto the same observation, implying non-uniqueness of the inverse problem (Baillet 2010).

*Components* (columns of **Y**) also belong to the subspace spanned by the columns *x_j_*(*t*) of **X**. Some analysis methods (e.g. PCA) result in columns of **Y** that are mutually uncorrelated, in which case the subspace spanned by any subset of columns is *orthogonal* to the subspace spanned by the other columns. In other words, each vector (signal) in one subspace is *uncorrelated* with every vector in the other subspace:

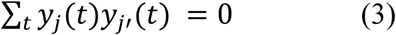

#### Spatial filter, null filter, and zero set

As mentioned earlier, a *spatial filter* is a vector of *J* weights [*u_j_*] to be applied to the observation channels. The source-to-filter-output gain for source *i* is:

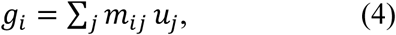

a weighted sum of the gains from that source to each sensor. A linear combination of filters is a filter, so spatial filters form a vector space, and because there are *J* coefficients this space is of dimension *J*.

A *null filter* **u** for a source is a filter that suppresses that source, i.e. the source-to-filter output gain is zero:

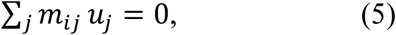

As long as there are at least two sensors (*J* > 1), a null filter for a source exists, a trivial example being a filter with all coefficients zero except for two channels *j* and *j*′ such that *u_j_*/*u_j’_* = − *m_ij_*/ *m_ij’_*. Since a weighted sum of null filters for a source is a null filter for that source, null filters for a source form a vector space, subspace of the space of spatial filters. Its dimension is *J* − 1 (the constraint of Eq. 5 reduces the dimension by one). In practical terms, every source has *many* null filters: many different linear combinations produce a signal that is “blind” to that source.

Given a second source *i*′, there exists likewise a subspace of null filters of dimension *J* − 1 for that second source. The null filters for *both sources* belong to the intersection of their subspaces, i.e., to a subspace of dimension *J* − 2. Repeating this argument, for *J* observation channels one can suppress up to *J* − 1 arbitrarily-positioned sources. The filter that does so belongs to a subspace of dimension one, i.e., the filter is *unique* to a multiplicative factor. For a greater number of sources, we cannot guarantee the existence of a filter that suppresses them all. For *J* channels, in general, *at most J* − 1 *sources can be suppressed by a spatial filter*.

Given a filter, the *zero set* of that filter is the set of source locations for which the gain (Eq. 4) is zero. If the filter is a null filter for a source, its zero set obviously includes that source. However, based on the reasoning that led to Fig. 3, we expect other sources to have the same gain (zero). In other words, we expect a zero set to be extended in source space, as will be apparent in several of the figures below.

#### Data-driven analysis

Spatial filters can be hard-wired as mentioned earlier, but it is not always easy to guess in advance the best *J* coefficients. Data-driven methods such as PCA, ICA, beamforming, or others offer a convenient way to find useful sets of coefficients. Among them, PCA stands out for three useful properties. First, the columns of the component matrix **Y** = **XU** are mutually uncorrelated (Eq. 3). Second, their variance adds up to the variance of **X**. Third, the first principal component (PC) packs the largest possible amount of variance, the second the largest amount of variance in the subspace orthogonal to the first, and so-on. A consequence of this third property is that the *last* principal component includes the *least* possible amount of variance and, indeed, if the data are rank-deficient its variance is *zero*. Thus, if at most *J* − 1 sources are active, the last column of the PCA matrix is a *null filter* for them all.

Other methods offer different properties (Parra et al 2015). ICA attempts to ensure that components are *statistically independent* according to some empirical measure (Cardoso 2001). Canonical correlation analysis (CCA) seeks linear combinations of columns of **X** that are *maximally correlated* with some other dataset **X**′. Joint decorrelation (JD, also known as denoising source separation, DSS, or common spatial patterns, CSP), seeks to maximize the *ratio of variance* between the data and a filtered version of the data (see de Cheveigné and Parra 2014 and references therein). *Beamforming* applied to a source location produces a filter that minimizes the variance at the output of the filter, subject that the gain from that location (Eq. 4) is one. Data-driven analysis is discussed further in a later section.

### Illustrations

#### A flat world

The previous concepts are somewhat abstract, and it may be hard to grasp the relations between *signal spaces*, the *physical space* that the sources and sensors inhabit, and the *parameter space* of a source model. In the real world, brain sources are typically assimilated to dipoles located within a 5D space (position and orientation) constrained to a convoluted shape (cortex), which is hard to visualize. The “flat world” captures the essence of the problem and is easier to visualize.

The flat world consists of a disk-shaped source space with sensors at the periphery (Fig. 4A), a geometry simpler than that of a brain surrounded by MEG sensors or EEG electrodes (panel B). Source-to-sensor gain varies with distance as 1⁄*d*^2^ and there is no dependency on orientation, in contrast to EEG or MEG. Panel C shows the gain pattern of a single sensor, and panel D shows the source-to-filter-output gain for a simple spatial filter with two non-zero coefficients, chosen to have zero gain for a source at the cross (this filter is a *null filter* for that source). The *zero set* of the filter in panel D includes the location of the source (cross), as well other locations along a curved line (white). In contrast to this simple picture, the gain patterns of a single MEG sensor (panel E) and a two-sensor spatial filter (panel F) are both convoluted, with bands of positive and negative gain according to relative orientations of sources and sensors. If source orientations were free (as opposed to normal to the cortical surface, as here), the 5D parameter space would be even harder to illustrate visually. The easier-to-visualize flat world captures the essence of the more complex system.

**Fig. 4.**
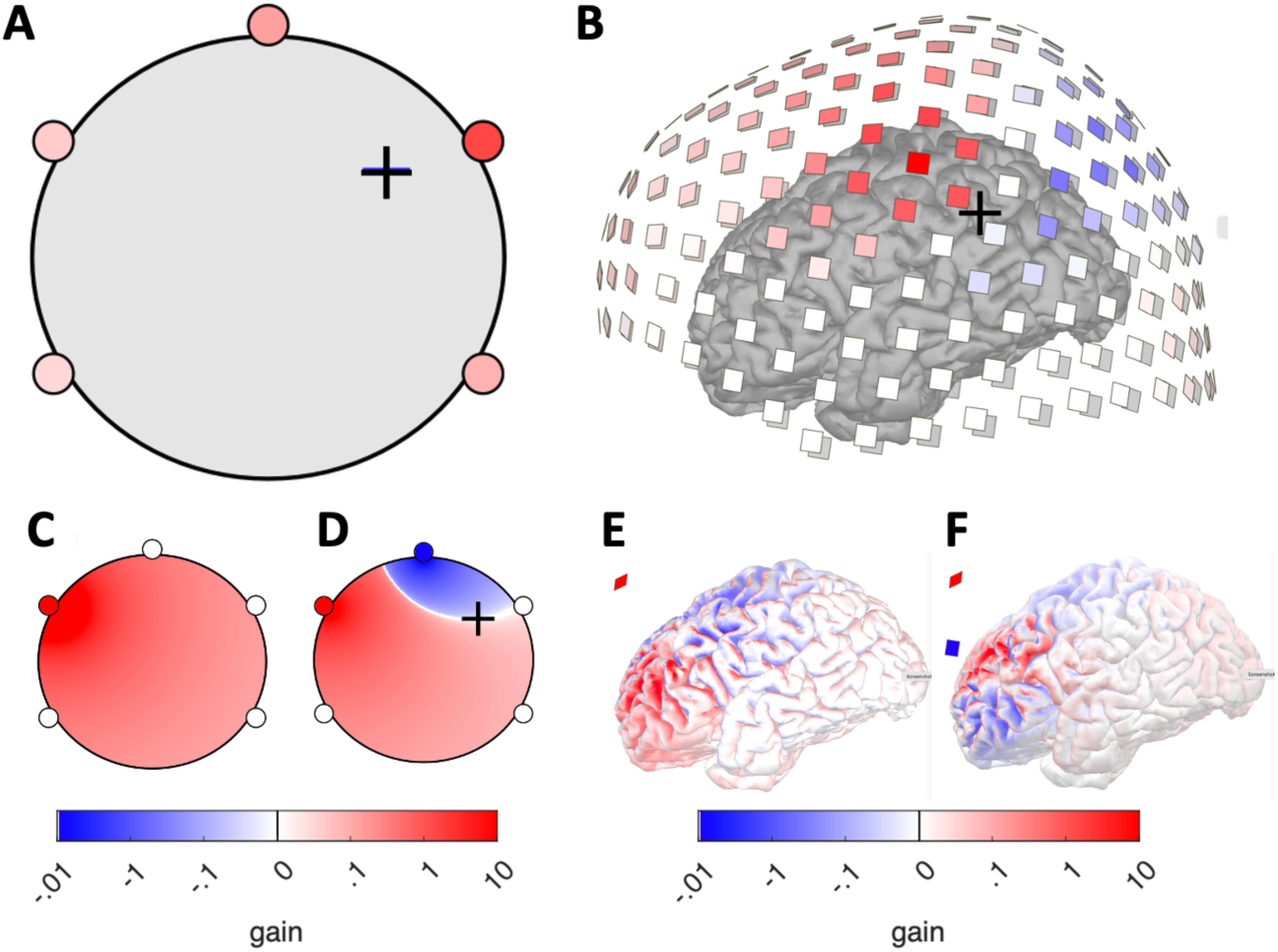
A: The “flat world” consists of a disk-shaped source space with sensors at the periphery. The source-to-sensor gain varies with distance as 1⁄*d*^2^ as represented by a color code at each sensor for a source located at the cross. B: The geometry of sources within the brain, and of sensors in a real MEG system are harder to visualize. The gain for each MEG sensor relative to a source is coded as color (the dipolar source is positioned at the cross, oriented normally with respect to the cortical surface, pointing inward). C: Gain pattern for a single sensor (red dot) as a function of source position in the flat world. D: Gain pattern for a simple spatial filter with two non-zero coefficients, chosen to ensure zero gain at the position of the cross. E: Gain pattern for single MEG sensor (oblique red square) relative to dipolar sources as a function of their location on the cortical surface. Sources are constrained to be normal to that surface. F: Gain pattern for a simple spatial filter with two non-zero coefficients with opposing sign (red and blue squares). All panels: positive and negative gains are plotted on separate logarithmic scales, with small and large absolute values clipped for visual clarity.

#### Null filters

Figure 5A shows the gain maps of four null filters for the source located at the cross. Their zero sets (white) have diverse shapes which all include the source location. These four null filters form a basis of the 4-dimensional space of null filters for this source, i.e. any null filter is a weighted sum of these (one example being the filter of Fig. 3D).

A word about how these filters were found: a Gaussian-distributed “source activity” matrix of size 1000×1 was multiplied by the 1×5 source-to sensor gain matrix for the location indicated by the cross, resulting in a “data matrix” of size 1000×5. PCA was applied, producing a 5× 5 transform matrix, the last four columns of which are null filters (Fig. 5A) for the source because the data matrix is of rank one. This illustrates the aforementioned process of deriving spatial filters by a data-driven method. For the 274-sensor MEG system of Fig. 3B there is likewise a 273-filter basis for the subspace of null filters for a source, but the gain pattern of that filter is harder to visualize (not shown).

**Fig. 5.**
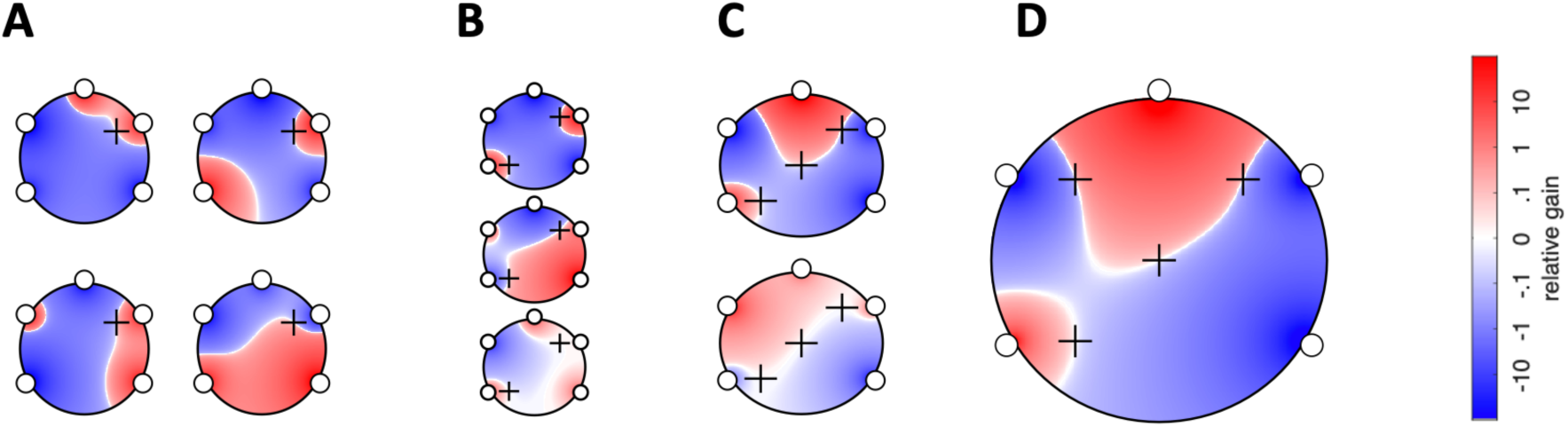
A: Gain maps for four null filters for the source located at the cross. B: Three null filters for two sources. C: Two null filters for three sources. D: Unique null filter for four sources.

#### Null filters for multiple sources

Figure 5B shows the gain maps of three null filters for *two* sources (crosses). Here, null filters form a subspace of dimension *J* − 2 = 3. Likewise, panels C and D show gain maps of null filters for three and four sources, respectively. The null filter for four sources (panel D) is *unique* to a multiplicative constant. That filter can suppress additional sources only if they fall within its zero set (white lines in panel D), otherwise, in general, at most *J* − 1 = 4 sources can be suppressed. For the 274-channel MEG system of Fig. 3B, at most 273 arbitrarily-located sources can be suppressed, and the filter that achieves this (without mapping everything to zero) is also *unique* to a multiplicative factor.

### A virtual electrode?

A real electrode has two properties that make it useful in scientific and clinical applications. First, the SNR (signal-to-noise ratio) of the activity of a neural source (e.g. single neuron or epileptic focus) can be enhanced by placing the tip near that source. Second, a source can be located by “listening” for some expected activity while moving the tip around. For example, a primary sensory region might be located by advancing an electrode until stimulus-evoked spikes are heard over the loudspeaker. Ideally, we’d like a virtual electrode to approximate both properties. I’ll look at each in turn.

#### SNR enhancement

The SNR is the ratio between the variance of the target (source of interest) and that of the background (all other sources). In a simulation where target and background are known, the SNR can be calculated exactly at each sensor and at the output of a spatial filter. We can then calculate the *SNR enhancement ratio* as the ratio between SNR at the filter output and the maximum SNR over sensors: a value greater than one indicates that the filter offers a benefit over just selecting the best sensor. If the number of background sources is *J* − 1 or less (as is the case for all filters illustrated in Fig. 5 and Fig. 6A), and the target source is not located within the zero set of the filter, the enhancement ratio is *infinite*. If the target is located within the zero set, the ratio is undefined (the filter suppresses everything). In the more general case of *J* or more background sources (as in real EEG or MEG recordings), the SNR enhancement ratio will depend on the geometry of all sensors and sources.

To illustrate the range of SNR enhancement ratios to expect, Fig. 6 shows gain maps for filters designed to optimize the SNR of a target (circle) in the presence of interferers (crosses), for various configurations. The filter in panel A achieves infinite enhancement because, with 15 sensors, the 14 interferers can be perfectly suppressed by a spatial filter, and the target luckily does not fall within its zero set. For panels B-E, the SNR ratio depends on the configuration in a complex fashion: compare C and D with same sensor and target layout but different interferer positions, and B and E with the same target and interferer layout but different sensor positions. The bottom line is (a) spatial filtering is always beneficial in terms of SNR, and (b) the benefit depends strongly on the configuration.

**Fig. 6.**
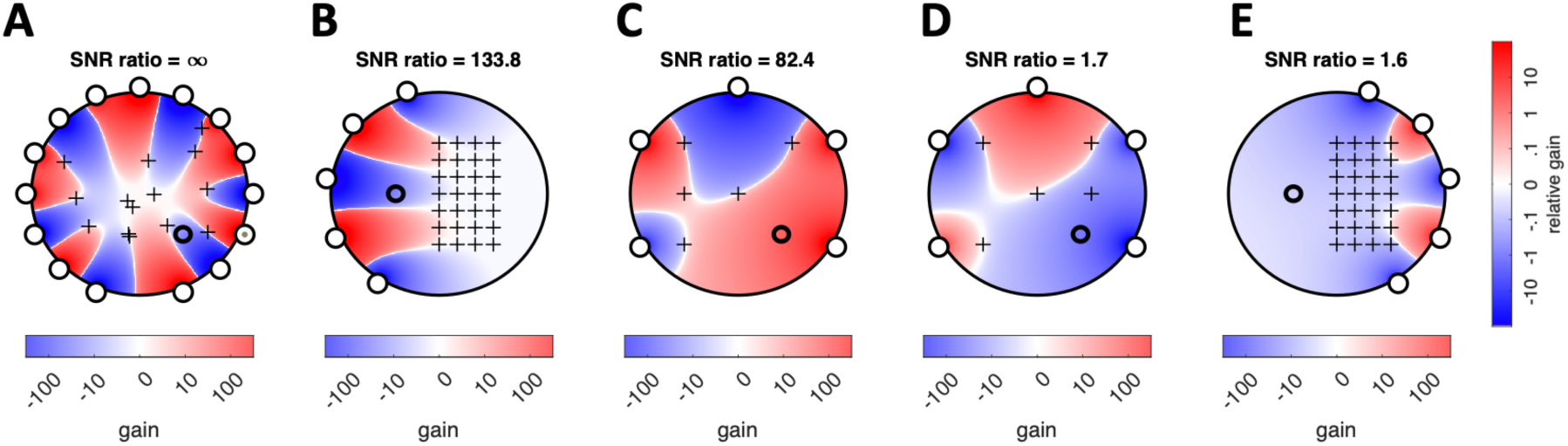
Quantifying SNR enhancement in a flat world model. A to E: Gain patterns for spatial filters that optimize the SNR of a target (circle) in the presence of background sources (crosses) for various layouts of target, background and sensors. The SNR enhancement ratio is indicated for each plot.

Figure 7 shows analogous results for a simulation with real MEG data and an anatomy-informed source/forward model based on publicly-available data from the BrainStorm tutorial. That dataset included 274-channel MEG recordings in response to repeated tones in an auditory oddball task, together with co-registered anatomical and sensor information. A gain matrix was calculated based on an overlapping sphere model, each column representing the gain from one source location to each of the sensors. The model sources were dipoles that could be located in any of 135002 positions on the cortical surface, with an orientation normal to the surface. A “target” source waveform was synthesized as a 1 Hz sinusoid with amplitude 30 nAm and multiplied, for each dipole location in the model, by the corresponding column of the gain matrix, and the result added to the real MEG data. The real MEG data (including eventual sensory and motor responses) were used only as a realistic background for this simulation.

For each source location, a LCMV beamformer solution was calculated to obtain an optimal filter to suppress the background while preserving the target. Figure 7A plots the maximum SNR over sensors (i.e. the best SNR achievable without spatial filtering) for each target location within the left hemisphere (viewed from the right, with the right hemisphere removed). As expected, the SNR tends to be better for superficial sources (red/yellow) than deep (blue). Panel B plots the SNR after spatial filtering: the SNR is improved for all target source locations, as confirmed by a plot of the SNR enhancement ratio (panel C). The enhancement ratio ranges across locations from ∼2 to ∼35 (median ∼5). Such values are of practical significance: an enhancement ratio of 5 is equivalent to multiplying by 25 the number of repeats in an evoked-response paradigm.

This is both good news, and bad. The good news is that there is a benefit for every location. The bad news is that the size of the benefit is smaller than expected from a real electrode. Also, it varies from one location to the next in a rather unpredictable fashion, reminiscent of the diversity observed in Fig. 6. On average, the SNR ratio is somewhat larger for superficial and deep sources than intermediate (Fig. 7D), but at every depth the spread is wide. The diversity likely reflects the fact that the zero set of the filter that attenuates the background has a complex geometry (as illustrated for the flat world in Fig. 6), and avoiding it is harder for some locations than others. Of course, the patterns of Fig. 7 are specific to the dataset chosen: for other datasets patterns might differ as a function of sensor configurations, subjects, or recordings.

**Fig. 7.**
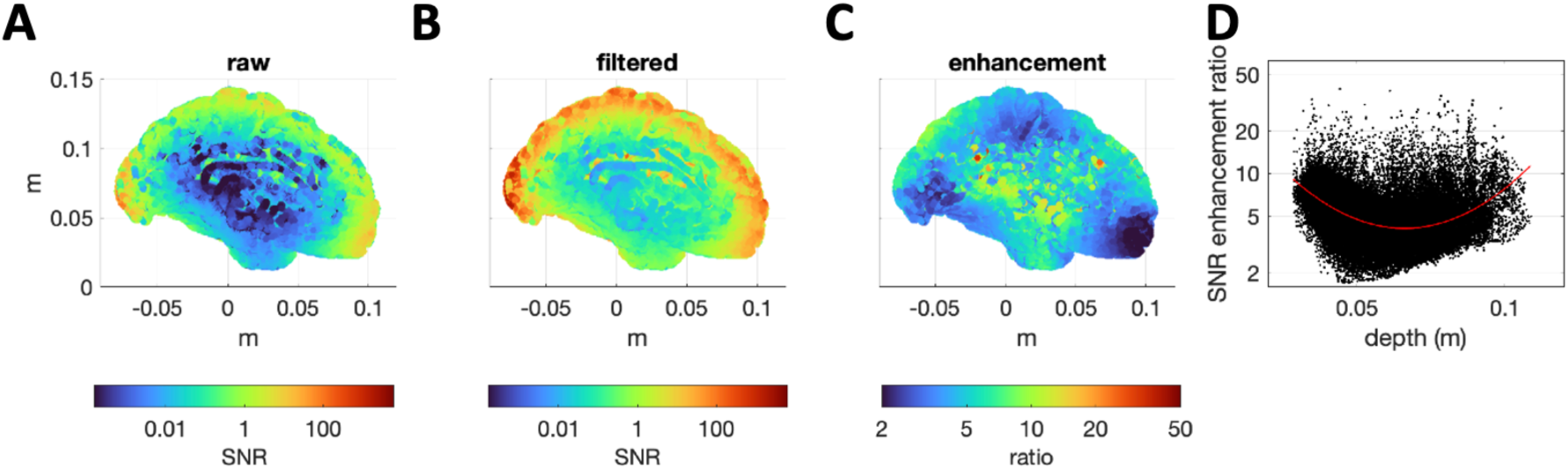
Quantifying SNR enhancement in a real 274-channel MEG configuration. A: SNR at the best sensor as a function of source location (left hemisphere viewed from the right, no transparency). B: SNR at the optimal filter output (same view). C: SNR enhancement ratio (same view). D: SNR enhancement as a function of depth (distance between a source and the closest sensor) for each source location. The red line indicates a quadratic fit to the log values.

The analysis illustrated in Fig. 7 might be useful to perform on a routine basis, so as to predict the benefit of a virtual electrode placed within a particular brain structure. It requires anatomical information (e.g. MRI scan), together with a recording representative of the expected background activity for that subject, but it does not require recording the target activity itself.

#### Source localization

A second useful property of a real electrode is the ability to locate a source by moving the electrode around until a particular response is observed. This property can be emulated by scanning a beamformer across the source space while monitoring its output. The location that yields the best SNR (according to some empirical measure) yields the position of the target source, as illustrated in Fig. 8A. The SNR is indeed maximal at the correct position (circle), but the pattern does not taper off with distance as fast as for a real electrode, and the peak is rather broad. Regardless, the principle works: a source can be located by scanning a model source space with a beamformer, as long there exists some criterion to detect when the filter output matches the target. Beamforming is one of several popular approaches to source localization (Baillet 2001, 2010; Jaiswal et al 2020).

It is interesting to observe that the filter found at the target location has the same gain pattern (panel B) as that of Fig. 5D, to a factor. Indeed, for *J* − 1 competing sources (as here), the filter found by the beamformer will be the same for every target source position (other than those that fall within the zero set of this filter). Thus, the gain pattern of this “virtual electrode” does not change as is moved around, counterintuitively and contrary to a real electrode. In the more general case of *J* or more competing sources, the gain pattern will change with target position, however we should not expect it to ever resemble the focal pattern of a real electrode.

**Fig. 8.**
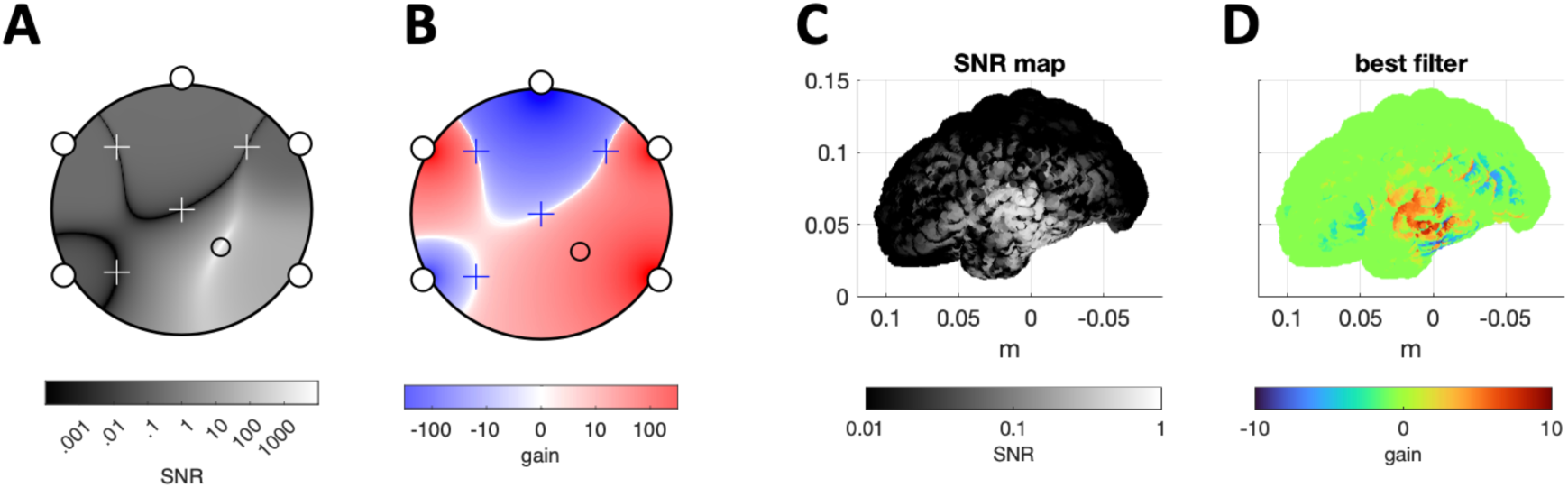
Locating a source by scanning with a beamformer. A: SNR as a function of the beamformer location parameter within the flat-world source space, for a target source (circle) in the presence of four competing sources (crosses). B: Gain map of the spatial filter found by the beamformer at the target location. C: SNR as a function of beamformer location parameter for a source located on the left-most node of a dipolar 274-channel MEG source model. D: Gain map of the beamformer solution when the its location parameter coincides with the target source. SNR plots are in gray because values are uniformly positive, gain plots are in color because values can be positive or negative (in contrast to Fig. 7 which used color for both for visual clarity).

Figure 8C shows the SNR of a spatial filter created by a beamformer as it is scanned across the parameter space of an MEG cortical source model. The data were created by placing a target source at the leftmost location in the source model (roughly centered on the white zone in panel C), applying the forward model for that position, and adding the result to a background of real MEG data. Analogous to panel A, the SNR is maximal at the correct position, but again the pattern does not taper off as fast as for a real electrode. The gain pattern of the filter obtained by the beamformer pointed at the target is shown in panel D. Again, it does not resemble the focal gain pattern of a real electrode.

### A virtual scalpel?

The ideal goal of a virtual electrode is incompletely attained, but an alternative metaphor worth exploring is that of a “virtual scalpel” by which the activity of a source is silenced within the data. This, we saw, can be achieved perfectly by a spatial filter (indeed the filters that do so form subspace of dimension *J* − 1 of the space of filters), suggesting an approach to analysis based on cancellation of individual sources. This can take several forms.

A first is *denoising*, by which an unwanted source is projected out of the data, for example by a denoising matrix defined as

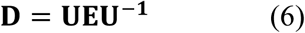

where **U**^-1^ is the inverse (or pseudoinverse) of the transform matrix **U**, and **E** is a diagonal matrix with ones for components to retain, and zeros for components to remove. Applying this to the data yields a “clean” data matrix **X̂** = **XD**. This matrix has the same size as the original data and can be substituted for it in further analyses, with two caveats. First **X̂** is not of full rank. Second, if further analyses call for a forward model, that model may need to be adjusted for effects of the denoising matrix (e.g. Hipp and Siegel 2015). An example of this approach is the common use of ICA to isolate and remove ocular artifacts, another is that of JD to remove eyeblinks or power line artifacts.

A second approach is illustrated in Fig. 9 for a flat-world simulation. Multiple spatio-temporal components are common for stimulus-evoked responses because latencies and/or time constants can differ between hemispheres and/or primary and higher-order fields, however it may be hard to separate them one from the other. The simulation includes two target sources (circles in lower parts of panels C-E) driven with distinct but correlated waveforms (panel A) that repeat periodically, and three competing sources (crosses in lower parts of panels C-E) driven with independent Gaussian noise. The sources were mixed with source-to-sensor gains appropriate for their positions, resulting in the mixture shown in panel B. A first JD analysis was applied to separate the targets from the noise, resulting in two components (panel C), both clean of noise but neither of which maps to a source (the corresponding filters have non-zero gain at both source locations). A second JD analysis was then applied to these two components to find an “early component” active from 1-100 samples. The first component of this analysis (panel D, blue) is again a mixture, but the second component (red) matches source 2 (indeed, the filter shown in the lower part of the panel cancels all but that source). A third JD analysis applied to find a “late” component (active from 201-400 samples) likewise found a component matching source 1. This approach is applicable to real EEG or MEG data (not shown), but the outcome is harder to validate for lack of ground truth.

**Fig. 9.**
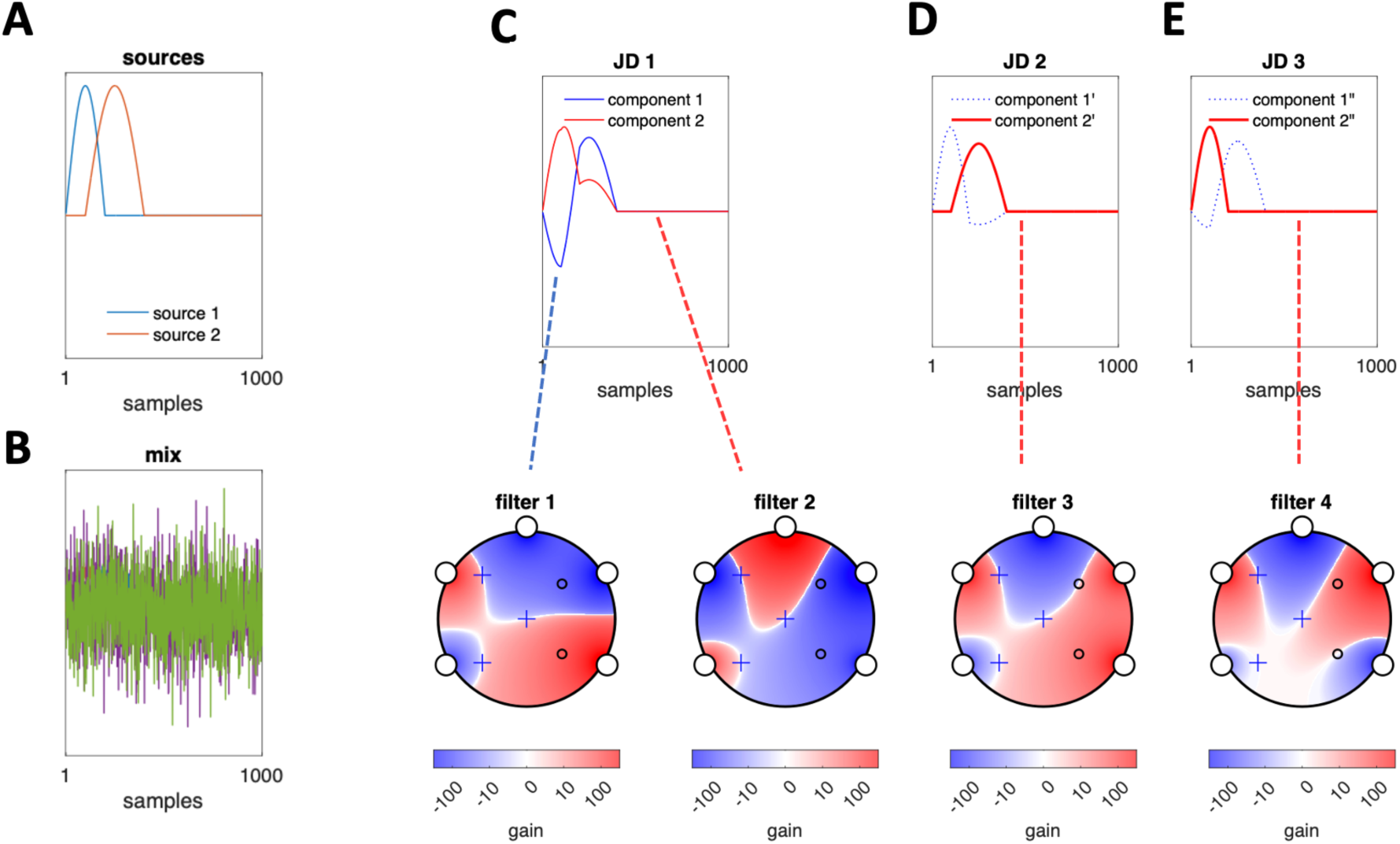
A: Time courses of target sources (with locations indicate as circles in panels C,D). B: mixture. C: Time courses of components retrieved by JD (top) and maps of filters associated with those components (bottom). D: Time courses of components rotated by JD to isolate the later source (top) and corresponding filter (bottom) E: Time courses of components rotated by JD to isolate the earlier source (top) and corresponding filter (bottom). The latter two filters (#3 and #4) have the property that they cancel all sources but one, which they thus isolate, a property that filters produced by the initial JD (#1 and #2) lack.

A third approach exploits the fact that a source is necessarily located at the intersection of the zero sets of all its null filters to localize it (de Cheveigné 2024, not discussed further here). These examples show how spatial filters might be used as sort of “virtual scalpel” to dissect complex brain activity. The first (denoising) is well established, the others are less thoroughly validated at this point.

### Finding a filter

Preceding sections focused on the properties of spatial filters; this section revisits how they are found. Trial-and-error is infeasible: trying just two values for each of 274 coefficients would require roughly as many trials as there are atoms in the universe (∼10^82^).

#### Hard-wired

Filter weights are chosen based on prior knowledge. For example, common-mode interference that affects all channels might be attenuated by subtracting one particular channel, or a combination of channels (re-referencing). If the interference is known to vary slowly across space (as for a distant environmental source in MEG), a differential montage may be useful (e.g. gradiometers in MEG). The spatial model can be more complex, as in the SSS method (Taulu and Simola 2006), implemented under the name of MaxFilter in certain MEG systems. The magnetic field is decomposed on a basis of spherical harmonics centered on the subject’s head, and this decomposition is used to determine weights to apply to the sensors that sample the field. The basis is partitioned into components likely to arise from within the head, which are retained, and those likely to arise from outside, which are discarded.

#### Data-driven

PCA involves eigen-decomposition of the empirical *covariance matrix* of the data (Greenacre et al 2022), while JD involves joint eigen-decomposition of two covariance matrices (de Cheveigné and Parra 2014 and references therein). Likewise, CCA exploits the covariance matrix of two concatenated datasets (de Cheveigné et al 2018 and references therein). Each of these methods “sees” the data through statistics that capture the correlation structure of the data (which itself reflects mixing of the same brain sources into multiple channels, Eq. 1), and ICA may additionally exploit higher-order statistics. None of these methods uses sensor location or anatomical information, and there is no assumption on the number or nature of the sources.

#### Hybrid (beamformer)

The LCMV beamformer solution for source location *i* is obtained as

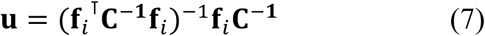

where **u** is the spatial filter, **f**_i_ is the source-to-sensor gain for that location, and **C** is, again, the covariance matrix of observed multichannel data (Van Veen et al 1997; Jaiswal et al 2020). Here, statistics from the data are complemented by a source/forward model which captures known geometry and anatomical constraints. The added constraint can be a plus (unrealistic solutions are avoided) or a minus (an optimal filter might not be found if the target does not fit the source model).

#### Existence vs identifiability

Data-driven analysis can fail for two reasons: (a) *the desired solution does not exist*, due to a fundamental constraint, (b) *the desired solution exists but we fail to find it*, for example due to geometrical inaccuracy, or lack of appropriate criteria, or overfitting. In the first case, a new technique might overcome the limitation; in the second, it cannot. It is important to distinguish these two situations.

#### Overfitting

Overfitting occurs when there are insufficient data to properly constrain the parameters. An oft-quoted rule of thumb is that at least 10*J* time points are needed for *J* parameters, but the theoretical basis for that rule is weak (Siegel 2022). In particular, overfitting may be *less severe* if inter-channel correlation allows dimensions to be reduced, or *more severe* if the data have serial correlation, as this implies fewer “effective samples.” Indeed, brain data often have roughly 1/*f* spectra and thus strong serial correlation. Overfitting entails both a risk of unsound observations (the analysis latches on to a quirk), and a loss of sensitivity in the cross-validation stage (the analysis fails to recognize a real effect). I recommend a three-pronged approach:

1. Proactively look for factors conducive to overfitting, and evidence that it might be occurring. Are the results too good to be true?
2. Deploy standard cross-validation techniques (e.g. Varoquaux 2017), such as training on a subset of the data and testing on another. Cross-validated results are less likely to be spurious.
3. Reduce dimensionality and/or apply regularization.

Cross-validation by itself does not “cure” overfitting. Rather, it reduces the risk of reporting a false result, but also makes analysis less sensitive to a real effect. Dimensionality reduction reduces the overfitting itself.

#### Is data-driven analysis safe?

Empirical sciences ask questions of the data, the foremost of which is whether an observed pattern is *real*, that is, it reflects an interesting phenomenon rather than a random quirk. A tool such as spatial filtering may help by reducing randomness and increasing the SNR of real patterns, however we must be worried when that tool itself depends on the data. For example, the JD algorithm is useful to find a spatial filter that isolates stimulus-evoked response components from multichannel neural data (de Cheveigné and Parra 2014). Applied to random data, it will oblige by discovering any chance directions that happen to repeat over trials. The investigator must be alert (if not paranoid) to such “double-dipping” (Kriegeskorte 2009; Gross et al 2013; Devezer et al 2021). Data-driven analysis is *not* safe: it requires attention.

#### What else can go wrong?

Disconcertingly, noise from one channel can be *injected* by a spatial filter into other channels. This is easily understood in the case of re-referencing: an artifact on the new reference (e.g. myogenic) will invade channels that were previously clean, with patterns that seem all the more “real” as they are now seen on all channels. However, the phenomenon can arise with any form of spatial filtering, often unexpectedly due to the hands-off nature of data-driven methods. There is no easy fix, other than be attentive to the phenomenon, and weigh the benefit of removing one source of noise against introducing another.

### Tips and tricks

#### In what order?

Spatial filtering is typically one step among several in a pipeline. Linear steps such as spatial or temporal filtering can be swapped, but non-linear steps (as involved in data-driven methods) cannot. For example, PCA followed by temporal filtering will not yield the same outcome as if temporal filtering comes first. Spatial filtering entails reducing the variance of background sources, but first removing some of that variance by other means (e.g. high-pass filtering) frees parameters to obtain a better spatial filter to remove the rest. Likewise, if high-amplitude glitches are removed first, the algorithm will not waste parameters trying to attenuate them. On the other hand, applying spatial filtering first might make those glitches easier to identify and remove. The order of operations counts, but there is no universal rule: in each situation a careful consideration of processing order may yield better results.

#### Stepwise dimensionality reduction

Dimensionality reduction reduces overfitting and speeds computation, however there is the risk of discarding informative dimensions by mistake. Furthermore, a data-driven dimensionality reduction operation may itself be prone to overfitting. A plausible approach is to operate in steps, each relatively conservative. For example, PCA can be applied to safely remove dimensions with variance below the floor of instrumental noise. It can be preceded by SCA (shared component analysis, de Cheveigné 2021) to down-weight variance specific to single channels, unlikely to reflect a neural source, and/or followed by JD to remove spatial components dominated by variance in an irrelevant spectral region, etc. For this to be of benefit, it is critical to verify that the criteria used to prioritize dimensions are distinct from effects tested in the study.

#### Boosting performance

A data-driven filter is the result of a compromise among various sources of variance to remove. The terms of this compromise can be improved in several ways. One is to first reduce the background variance by other means, as mentioned above. As an example, repeating the simulation of Fig. 7 with a 30 Hz low-pass filter applied to the MEG background would increase the median SNR enhancement ratio by a factor of two, which, added to direct effects of low-pass filtering, would boost SNR by a factor close to three. Thus, if activity above 30 Hz is not of interest, initial low-pass filtering is a good move. A second way is to increase the dimensionality *J* of the data. Adding more sensors increases the number of sources that can be suppressed, with two caveats (in addition to cost): each sensor contributes its own noise, adding to the count of sources to be cancelled, and it also increases risks of overfitting. If these factors are controlled for, increasing the number of sensors or electrodes is beneficial.

Data dimensionality can be increased in other ways. Taking advantage of the temporal *sparsity* of many sources, a different filter may be applied to different partitions of the time axis. With *N* intervals, the effective dimensionality jumps to *NJ*. Alternatively, the data may be filtered into different frequency regions, each with a different spatial filter. This approach can be generalized by augmenting the data with time shifts [ 0 ⋯ *N* − 1] (or some other convolutional basis): the spatial filter is then replaced by a *NJ*-coefficient *spatio-temporal* filter (multichannel finite-impulse-response filter). Yet another approach is to operate within a space of quadratic forms based on cross-products of the form *x_j_*(*t*)*x_j_*_’_(*t*). These can then be filtered by a *J*(*J* − 1)/2-coefficient *quadratic component analysis* filter (QCA, de Cheveigné 2012), sensitive to power.

Each of these schemes entails increased risks of overfitting, the burden of mastering a more complex technique and explaining it to the reader, and time wasted “tinkering” with its various possibilities. It also increases *researcher degrees of freedom*, which should not be a deterrent but requires care when evaluating statistical significance (Kriegeskorte et al 2009; Simmons et al 2021).

### Spatial filtering “under the hood”

There may be contexts where it is not obvious that spatial filtering is involved. For example, “PCA (or beamforming, etc.) was applied to the data…” should be read as “the data were spatially filtered with a filter found by PCA (or beamforming, etc.)…”. A decoding model or a brain-computer interface (BCI) may include an initial spatial filtering stage, the role of which is, implicitly, to unravel the linear mixing that produced the data. Neural networks are likely to have an initial linear layer for the same reason, and it is worth noting that the popular RELU (rectified linear unit) is piecewise linear: bias from another node can shift a unit between linear regimes, thus switching between spatial filters. Realizing that spatial filtering is involved may help to understand the properties of these methods and aid in their design. In particular, neural networks can benefit from prior knowledge (“inductive bias”) to avoid overfitting.

## Conclusion

Spatial filtering is an important tool for brain data analysis, that goes some way towards reversing the source-to-sensor mixing process that produced the data. In this, it is limited by the sparsity of observations that implies a non-uniqueness of the inverse problem (Baillet 2001, 2010). A well-chosen spatial filter can increase the SNR of sources of interest beyond that available at any of the sensors, which motivates the term “virtual electrode” to describe such a filter. It may be tempting to over-state this metaphor, and consider non-invasive exploration as an alternative to invasive exploration with a real electrode, however there are limits to the ability of spatial filtering to fill this role.

What a spatial filter can do perfectly is *cancel* a source, and indeed, with *J* sensors it can cancel up to *J* − 1 sources. The required filter exists, however (a) if there are other interfering sources, it may not suffice to elevate the SNR of a target brain activity to a useful level, and (b) it may not be possible to find it for lack of an appropriate criterion to support data-driven filter estimation, or because of overfitting. A larger number of channels allows more sources to be cancelled, but may increase overfitting.

A goal of this paper was to ease understanding of these factors, and the algebraic, geometrical, physical and computational constraints behind them, which are made obscure by the complex geometry of brain sources and sensors, and the high dimensionality of the data.

## Acknowledgments

Malcolm Slaney, Lucas Parra and Israel Nelken and two anonymous reviewers gave useful comments on earlier drafts. This work was supported by grant ANR-17-EURE-0017.

## References

Alberto, G. E., Stapleton-Kotloski, J. R., Klorig, D. C., Rogers, E. R., Constantinidis, C., Daunais, J. B., & Godwin, D. W. (2021). MEG source imaging detects optogenetically-induced activity in cortical and subcortical networks. Nature Communications, 12(1), 5259. 10.1038/s41467-021-25481-y

Baillet, S., Mosher, J.C., and Leahy, R.M. (2001) Electromagnetic brain mapping. IEEE Signal Processing Magazine, 18(6):14–30.

Baillet, S. (2010) The Dowser in the Fields: Searching for MEG Sources, in MEG, an Introduction to Methods, edited by Hansen, P.C., Kringelbach, M.L., Salmelin, R., Oxford University Press, 83–123.

Cao, M., Galvis, D., Vogrin, S. J., Woods, W. P., Vogrin, S., Wang, F., Woldman, W., Terry, J. R., Peterson, A., Plummer, C., & Cook, M. J. (2022). Virtual intracranial EEG signals reconstructed from MEG with potential for epilepsy surgery. Nature Communications, 13(1), 994. 10.1038/s41467-022-28640-x

Cardoso, J.F., 2001. The three easy routes to independent component analysis; contrasts and geometry. Proc. Int. Conf. on Independent Component Analysis and Blind, Source Separation (ICA01), pp. 1–6.

Coelli, S., Medina Villalon, S., Bonini, F., Velmurugan, J., López-Madrona, V. J., Carron, R., Bartolomei, F., Badier, J.-M., & Bénar, C.-G. (2023). Comparison of beamformer and ICA for dynamic connectivity analysis: A simultaneous MEG-SEEG study. NeuroImage, 265, 119806. 10.1016/j.neuroimage.2022.119806

de Cheveigné, A. (2012) Quadratic component analysis, NeuroImage, 59, 3838–3844, 10.1016/j.neuroimage.2011.10.084.

de Cheveigné, A., Parra, L. (2014) Joint decorrelation: a versatile tool for multichannel data analysis. NeuroImage, 98, 487–505, doi: 10.1016/j.neuroimage.2014.05.068.

de Cheveigné A, Wong DDE, Di Liberto GM, Hjortkjaer J, Slaney M, Lalor E (2018) Decoding the auditory brain with canonical correlation analysis. NeuroImage 172, 206– 216, 10.1016/j.neuroimage.2018.01.033.

de Cheveigné, A (2021). Shared Component Analysis. NeuroImage, 226, 117614, 10.1016/j.neuroimage.2020.117614.

de Cheveigné, A. (2024) What and Where in Electromagnetic Brain Imaging (2024), *bioRxiv*, https://www.biorxiv.org/content/10.1101/2024.04.17.589904.

Devezer, B, Navarro, D.J., Vandekerckhove, J., & Ozge Buzbas, E. (2021). The case for formal methodology in scientific reform. Royal Society Open Science, 8, 200805.

Greenacre, M., Groenen, P. J. F., Hastie, T., D’Enza, A. I., Markos, A., & Tuzhilina, E. (2022). Principal component analysis. Nature Reviews Methods Primers, 2(1), 100. 10.1038/s43586-022-00184-w

Gross, J., Baillet, S., Barnes, G. R., Henson, R. N., Hillebrand, A., Jensen, O., Jerbi, K., Litvak, V., Maess, B., Oostenveld, R., Parkkonen, L., Taylor, J. R., Van Wassenhove, V., Wibral, M., & Schoffelen, J.-M. (2013). Good practice for conducting and reporting MEG research. NeuroImage, 65, 349–363. 10.1016/j.neuroimage.2012.10.001

Helmholtz H. (1867). Handbuch der Physiologischen Optik (English tranl.: 1924 JPC Southall as Treatise on Physiological Optics) Voss.

Hipp, J. F., & Siegel, M. (2015). Accounting for Linear Transformations of EEG and MEG Data in Source Analysis. PLOS ONE, 10(4), e0121048. 10.1371/journal.pone.0121048

Jaiswal, A., Nenonen, J., Stenroos, M., Gramfort, A., Dalal, S. S., Westner, B. U., Litvak, V., Mosher, J. C., Schoffelen, J.-M., Witton, C., Oostenveld, R., & Parkkonen, L. (2020). Comparison of beamformer implementations for MEG source localization. NeuroImage, 216, 116797. 10.1016/j.neuroimage.2020.116797

Juárez-Martinez, E. L., Nissen, I. A., Idema, S., Velis, D. N., Hillebrand, A., Stam, C. J., & van Straaten, E. C. W. (2018). Virtual localization of the seizure onset zone: Using non-invasive MEG virtual electrodes at stereo-EEG electrode locations in refractory epilepsy patients. NeuroImage: Clinical, 19, 758–766. 10.1016/j.nicl.2018.06.001

Kriegeskorte, N., Simmons, W. K., Bellgowan, P. S. F., & Baker, C. I. (2009). Circular analysis in systems neuroscience: The dangers of double dipping. Nature Neuroscience, 12(5), 535–540. 10.1038/nn.2303

Parra, L.C., Spence, C.D., Gerson, A.D., Sajda, P., 2005. Recipes for the linear analysis of EEG. Neuroimage 28, 326–341.

Simmons, J. P., Nelson, L. D., & Simonsohn, U. (2021). False-Positive Psychology: Undisclosed Flexibility in Data Collection and Analysis Allows Presenting Anything as Significant. Psychological Science, 22, 1359–1366.

Tadel, F., Baillet, S., Mosher, J. C., Pantazis, D., & Leahy, R. M. (2011). Brainstorm: A User-Friendly Application for MEG/EEG Analysis. Computational Intelligence and Neuroscience, 2011, 1–13. 10.1155/2011/879716

Tarantola, A. (2006). Popper, Bayes and the inverse problem. Nature Physics, 2(8), 492–494. 10.1038/nphys375

Taulu, S., & Simola, J. (2006). Spatiotemporal signal space separation method for rejecting nearby interference in MEG measurements. Physics in Medicine and Biology, 51(7), 1759–1768. 10.1088/0031-9155/51/7/008

Varoquaux, G., Raamana, P. R., Engemann, D. A., Hoyos-Idrobo, A., Schwartz, Y., & Thirion, B. (2017). Assessing and tuning brain decoders: Cross-validation, caveats, and guidelines. NeuroImage, 145, 166–179. 10.1016/j.neuroimage.2016.10.038

Van Beest, E. H., Bimbard, C., Fabre, J. M. J., Dodgson, S. W., Takács, F., Coen, P., Lebedeva, A., Harris, K. D., & Carandini, M. (2025). Tracking neurons across days with high-density probes. Nature Methods, 22(4), 778–787. 10.1038/s41592-024-02440-1

Van Veen, B. D., Van Drongelen, W., Yuchtman, M., & Suzuki, A. (1997). Localization of brain electrical activity via linearly constrained minimum variance spatial filtering. IEEE Transactions on Biomedical Engineering, 44(9), 867–880. 10.1109/10.623056

Velmurugan, J., Badier, J.-M., Pizzo, F., Medina Villalon, S., Papageorgakis, C., López-Madrona, V., Jegou, A., Carron, R., Bartolomei, F., & Bénar, C.-G. (2022). Virtual MEG sensors based on beamformer and independent component analysis can reconstruct epileptic activity as measured on simultaneous intracerebral recordings. NeuroImage, 264, 119681. 10.1016/j.neuroimage.2022.119681

Vrba, J., & Robinson, S. E. (2001). Signal Processing in Magnetoencephalography. Methods, 25(2), 249–271. 10.1006/meth.2001.1238

Wójcik, D.K. (2022). Current Source Density (CSD) Analysis. In: Jaeger, D., Jung, R. (eds) Encyclopedia of Computational Neuroscience. Springer, New York, NY. 10.1007/978-1-0716-1006-0_544

